# Isoprene deters insect herbivory by priming plant hormone responses

**DOI:** 10.1101/2024.11.01.621578

**Authors:** Abira Sahu, Mohammad Golam Mostofa, Yuan Xu, Bianca M. Serda, James O’Keefe, Thomas D. Sharkey

## Abstract

Isoprene, emitted by some plants, enhances plant abiotic resilience but its role in biotic stress resilience remains elusive. We used tobacco plants engineered to emit isoprene (IE) and the corresponding azygous non-emitting control (NE) to investigate isoprene emission and biotic stress resilience. IE plants were more resistant to insect herbivory than NE plants. Worms preferred to feed on NE rather than IE leaves. IE plants showed less decline in photosynthesis during worm feeding. Insect feeding increased jasmonate levels in IE leaves, suggesting isoprene-mediated priming of the jasmonic acid response. Wound-induced increase in isoprene emission corresponded with elevation of methyl-D-erythritol-4-phosphate pathway and Calvin-Benson cycle metabolites. The results highlight interactive functions of isoprene and jasmonic acid and advance our understanding of how isoprene emission enhances plant resilience.

## Introduction

Plants encounter a variety of environmental stresses throughout their life cycle, with pest infestations being one of the common challenges. Annual crop loss to pest attacks ranges between 20-40% (*1*), resulting in a global economic loss of $70 billion. The yield reduction is further exacerbated by global warming since elevated temperatures accelerate insect metabolism, leading to increased consumption of plant tissues (*2, 3*). Global yield loss to insects is anticipated to increase by 10 to 25% for each degree temperature rise (*4*). Pest attacks have a negative impact on several aspects of food security and economic viability including production costs and stability, distribution efficiency, crop nutritional value, and biomass quality (*5*). Hence, increased pest infestation necessitates greater pesticide application to mitigate yield loss, further deteriorating plant health and environment. Therefore, it is crucial to find or develop plants with enhanced pest resistance for ensuring global food security and economic viability.

Synthesis and emission of volatile organic compounds (VOCs), particularly terpenes such as monoterpenes and sesquiterpenes, is a key defense strategy employed by plants to counteract insect herbivory (*6-9*). In addition to these, many plants emit a hemiterpene, isoprene, which accounts for 50-60% of the total global non-methane hydrocarbon emission from the biosphere (*10*). Although isoprene emission can contribute to ozone and aerosol formation (*11, 12*), it plays a crucial role in protecting plants from abiotic stresses including high temperature, drought stress, and ozone stress (*13-20*). Isoprene synthesis through the methyl-D-erythritol-4-phosphate (MEP) pathway requires 14 NADPH and 20 ATP molecules, making it metabolically expensive (*21*). This metabolic cost is presumably outweighed by its benefits. Indeed, isoprene acts as a signaling molecule, modulating the plant transcriptome, proteome, and metabolome to enhance resilience against various environmental stresses (*22-24*). However, the role of isoprene in biotic stress tolerance and the associated underlying mechanistic pathways are largely unexplored.

Plants have evolved complex signaling mechanisms involving phytohormones, particularly jasmonic acid (JA), to defend against insect herbivory (*25*). Within 2 h of pest infestation, there is a rapid increase in endogenous JA and its bioactive derivative, jasmonoyl isoleucine (JA-Ile) (*26*). Additionally, JA-deficient mutants are more susceptible to insect feeding than wild-type (WT) plants, highlighting the crucial role of JA in insect resistance (*27, 28*).

Isoprene modulates the expression of several JA-biosynthetic genes in unstressed plants (*22*). Additionally, most isoprene-responsive genes contain several *cis*-regulatory elements that interact with transcription factors involved in JA signaling (*29*). Furthermore, genes involved in biosynthesis of stress-related metabolites including glucosinolates, polyamine, oxylipin, apiose, and phenylpropanoids are upregulated in isoprene-emitting plants (*22*). Isoprene also promotes phosphorylation of NO_3_-induced (NOI) peptides (*23*). Genes of the NOI family are upregulated in *Arabidopsis* plants after spider mite feeding (*30*). These results indicate the potential functions of isoprene in eliciting plant defense responses against insect herbivory. Tobacco hornworms (*Manduca sexta*) preferred feeding on non-emitting plants over isoprene-emitting plants (*31*), but there is a gap in our understanding of the mechanism(s) underlying isoprene-mediated plant defense responses to insect herbivory.

In this study, we evaluated the impact of isoprene emission on insect herbivory using homozygous isoprene-emitting (IE) tobacco plants transformed with *Populus alba isoprene synthase* (*ISPS*) and the corresponding azygous (NE) control. We identified the deterring effect of isoprene on whitefly infestation and hornworm development in IE tobacco plants. We assessed the changes in the hormone profile in IE leaves following insect feeding and wounding. We also evaluated wound-induced isoprene emission and its connection with MEP pathway and Calvin-Benson cycle (CBC) metabolites in IE plants.

## Results

### Isoprene deters whiteflies infestation and hornworm herbivory

To assess the impact of isoprene emission on plant resistance to insect attack, transgenic IE tobacco plants (hereafter referred as IE plants) were grown alongside NE plants in the same pot. Insect infestation was quantified by counting the number of whiteflies per cm^2^ leaf area of 8-w-old IE and NE plants (Fig. 1A). The number of whiteflies was significantly lower in IE plants than NE plants (Fig. 1B). To further evaluate the impact of isoprene emission on insect herbivory, an equal number of tobacco hornworm 1^st^ instar larvae were placed on IE and NE plants and leaf area consumption was quantified after 10 d of worm feeding. The hornworms ate less of the leaves from the IE plants compared to the NE plants (Fig. 2A). The hornworms fed on IE leaves exhibited growth reduction, leading to significantly lower larval weight compared to those feeding on NE leaves (Fig. 2B,C).

**Fig 1.**
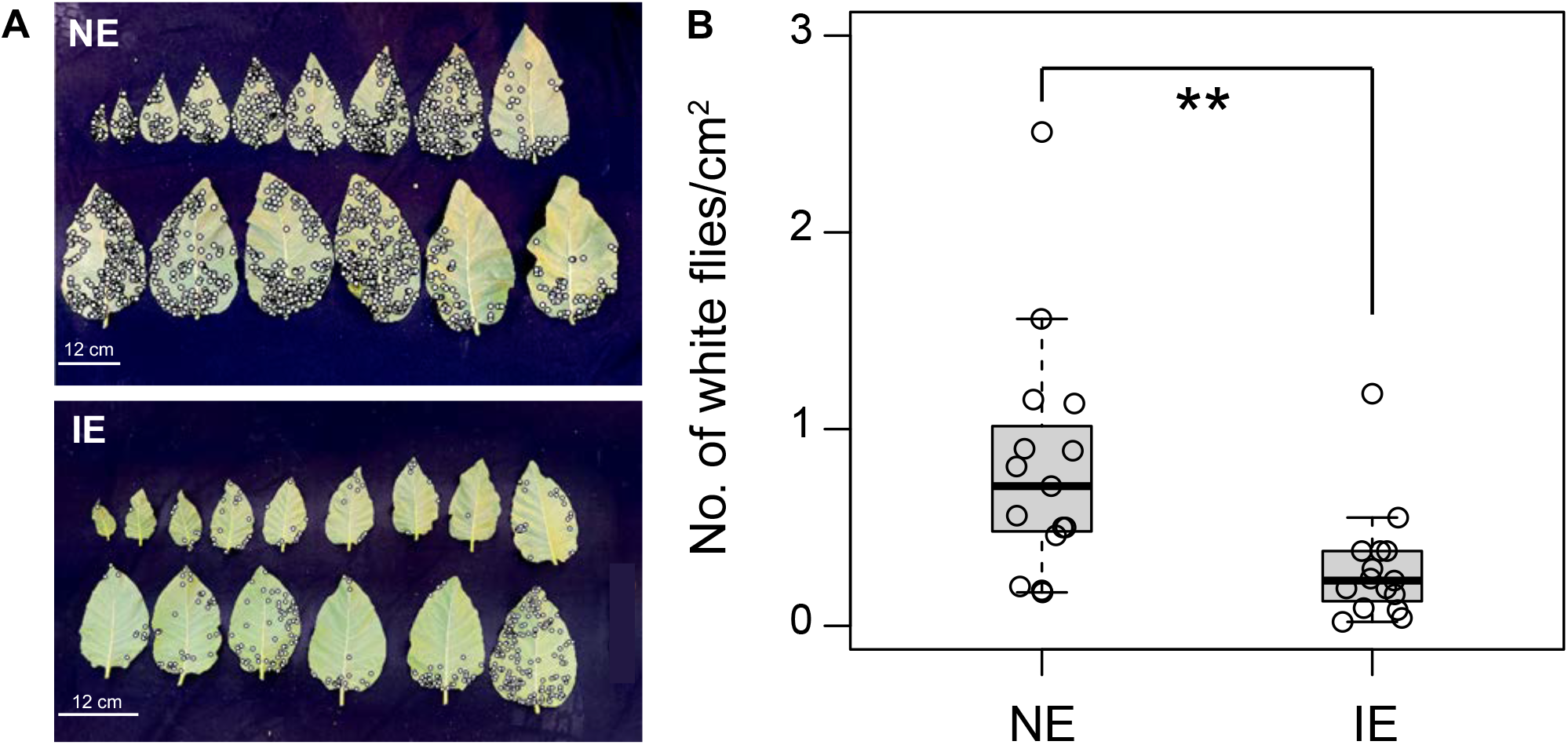
Effect of isoprene emission on white fly infestation. **(A)** White flies (shown by white dots) in isoprene-emitting (IE) and non-emitting (NE) tobacco plants grown together in the same pot (*n*=12-15 leaves). **(B)** Quantification of white fly infestation per cm^2^ leaf area in NE and IE plants. Asterisks indicate significant decline in whitefly infestation in IE leaves compared to NE leaves (*P*<0.01; Student’s t-test). Whiskers of the box plots represent 95% confidence interval.

**Fig 2.**
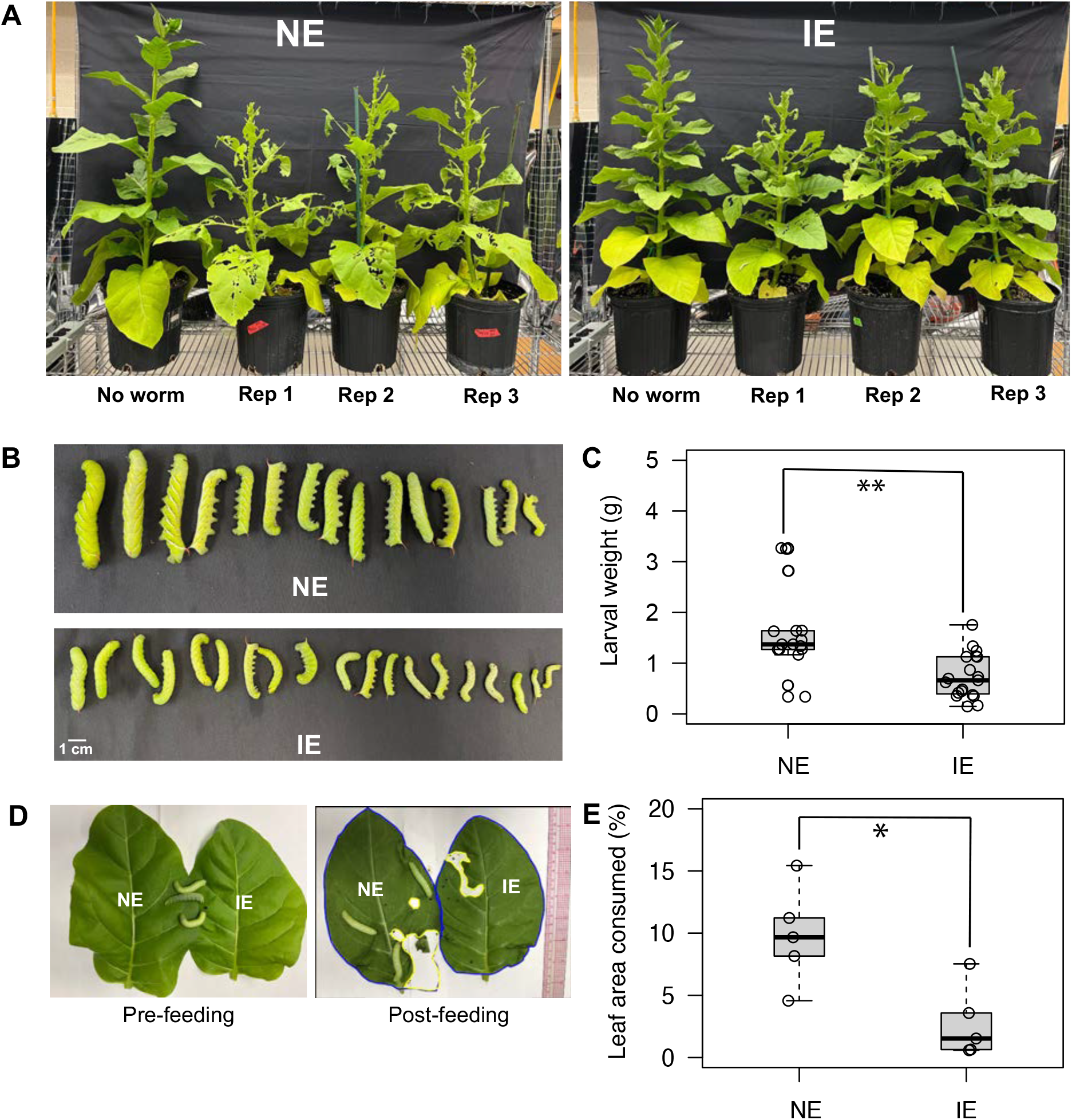
Effect of isoprene emission on insect herbivory and hornworm feeding preference study. **(A)** Comparison of leaf consumption by tobacco hornworms in NE and IE plants (*n*=3). (**B-C**) Comparison of hornworm larval weight reared on IE and NE plants for 10 days (*n*=15-18). Asterisks indicate significantly lower leaf consumption in IE leaves compared to NE leaves (*P*<0.01; Student’s t-test). **(D)** Comparison of leaf consumption by 10-d old worms when given the choice between NE and IE leaves. **(E)** Quantification of leaf consumption 90 min after placing worms on a pair of IE and NE leaf (*n*=4). Asterisk indicates significantly lower weight of worms feeding on IE leaves compared to NE leaves (*P*<0.05; Student’s t-test). Whiskers of the box plots represent 95% confidence interval.

### Isoprene influences hornworm feeding preference

An *in vivo* feeding preference study was conducted by placing 3^rd^ instar hornworm larvae on a pair of IE and NE leaves (Fig. 2D). After 90 min of feeding, the hornworms consumed significantly less amount of the IE leaves compared to the NE leaves (Fig. 2E). Real-time video monitoring revealed that, although the worms crawled over the IE leaf, they refrained from feeding on it and preferred to feed on the NE leaf instead (Movie S1). Upon closely reviewing the video, we noticed that one worm spent 26% of the recorded time on the IE leaf, however, it did not consume any part of the leaf. After staying on the IE leaf for 7-8 min, the hornworm eventually moved towards the NE leaf, and began to feed on it. To further evaluate their feeding preference, 1^st^ instar hornworm larvae were placed in a box containing a pair of IE and NE leaves (Fig. S1A). After 7 d of feeding, consumption of the IE leaf was significantly lower compared to the NE leaf (Fig. S1B). Additionally, worms recovered from the IE leaf after 7 d exhibited significantly lower larval weight compared to those on the NE leaf (Fig. S1C).

### Change in photosynthetic biochemistry in NE and IE leaves during worm feeding

To investigate the effect of insect herbivory on photosynthesis, gas exchange features including assimilation rate (*A*), intercellular CO_2_ concentration (*C*_*i*_), and stomatal conductance (*g*_*sw*_) were recorded in NE and IE leaves during worm feeding. Insect feeding induced stomatal closure (lower *g*_*sw*_), resulting in decreased *C*_*i*_ and *A* in the undamaged areas of both IE and NE leaves (Fig. 3A-C, S2A-C). However, IE leaves showed significantly lower reduction in *A* than NE leaves (Fig. 3D). The milder effect of herbivory on *A* was consistent with a smaller decrease in *C*_*i*_ in worm-fed IE and NE leaves (Fig. 3E, F). *A/C*_*i*_ response curves revealed that this distinct photosynthetic response in IE leaves was unrelated to differences in rubisco activity, electron transport, and triosephosphate use limitation since these parameters were similar between NE and IE leaves (Fig. 3G, Table S1), further supporting stomata-mediated photosynthesis decline.

**Fig 3.**
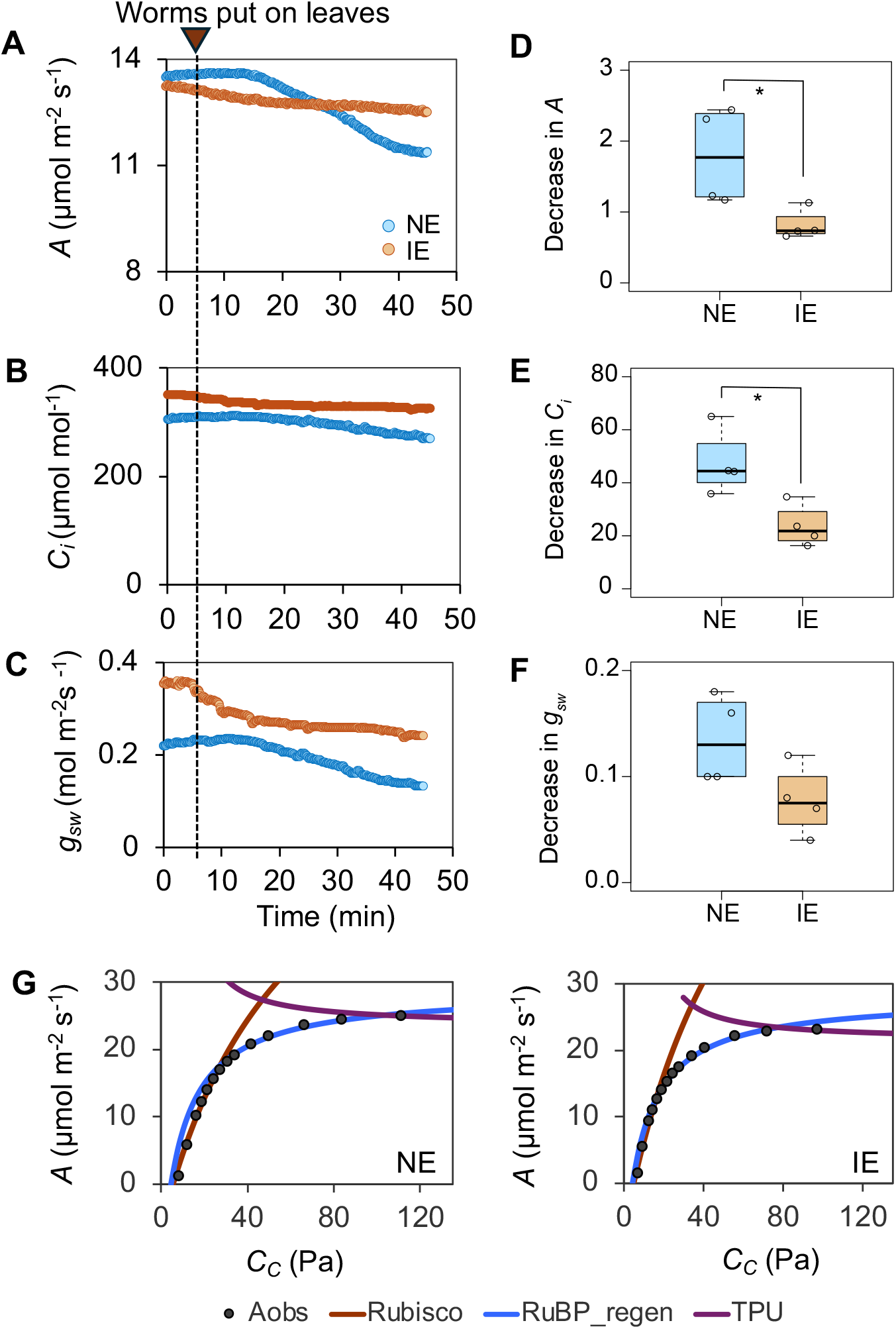
Effect of worm feeding on photosynthesis (*A*), intercellular CO_2_ concentration (*C*_*i*_), and stomatal conductance (*g*_*sw*_) in NE and IE leaves. **(A-C)** Changes in *A, C*_*i*_, and *g*_*sw*_ recorded during worm feeding in NE and IE leaves. **(D-F)** Comparison of absolute changes in *A, C*_*i*_, and *g*_*sw*_ pre-and post-feeding in NE and IE leaves. Decline in *A* and *C*_*i*_ is significantly lower in IE leaves compared to NE leaves (*P*<0.05; Student’s t-test). Whiskers of the box plots represent 95% confidence interval. **(G)** Representative *A/C*_*i*_ curves made using the dynamic assimilation technique in NE and IE leaves during worm feeding. Black filled circles are the measured CO_2_ assimilation rates, the red line represents the fitted rubisco *V*_*cmax*_, blue line is the fitted *J*, and purple line shows the triosephosphate use (TPU) limitation. No TPU limitation was observed during worm feeding.

### Hornworm feeding triggers differential responses in phytohormones in NE and IE plants

Levels of JA, JA-Ile, salicylic acid (SA), salicylic acid glucoside (SAG), and abscisic acid (ABA) were quantified in IE and NE leaf samples collected after 1 h of worm feeding. While undetectable in pre-feeding leaves, the JA level was significantly elevated in the leaves of both IE and NE plants post-feeding. Notably, the increase in JA in IE leaves was significantly higher than NE leaves (Fig. 4A). JA-Ile showed a similar trend, although the increase was not statistically significant (*P* = 0.06; Fig. 4B). SA was not detected in any sample, both pre-and post-feeding. Moreover, there were no significant differences in SAG and ABA levels between NE and IE leaves before and after hornworm feeding (Fig. 4C, D). To determine if this hormone response persisted over the long-term, leaf samples were collected from NE and IE plants 10 d after insect feeding. For control, leaf samples were collected from IE and NE plants of the same age that were never exposed to hornworms. Notably, all the studied hormones were similar after long-term herbivory, except for ABA. The ABA level was significantly higher in hornworm-fed IE leaves (Fig. S3).

**Fig 4.**
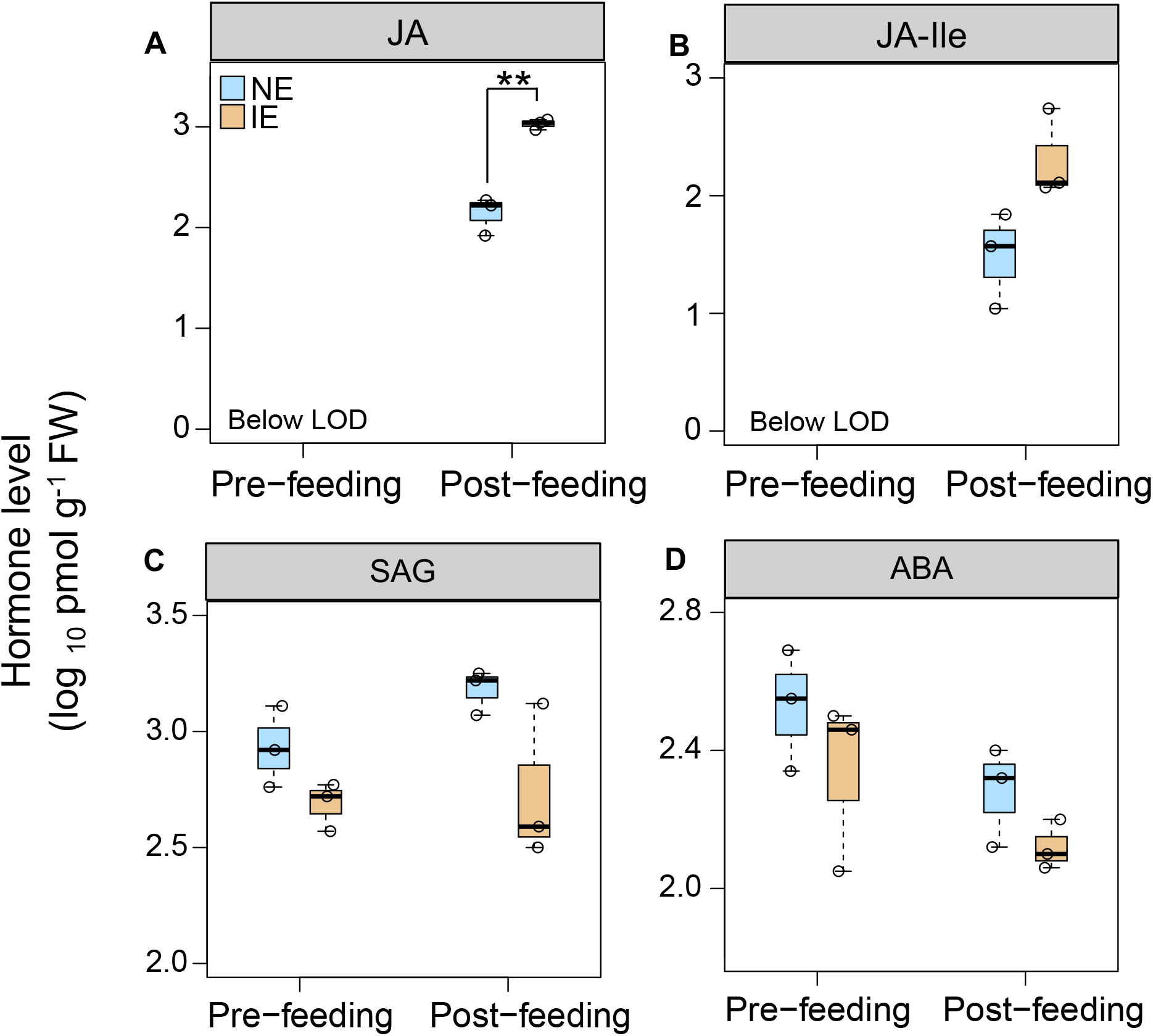
Change in hormone levels in NE and IE leaves after worm feeding. Hormones were quantified in leaves 1 h post-feeding (*n*=3). **(A)** Post-feeding JA level was significantly higher in IE leaves compared to NE leaves (*P*<0.01; Student’s t-test). Change in **(B)** JA-Ile, **(C)** SAG, and **(D)** ABA levels between NE and IE lines was not statistically significant. Whiskers of the box plots represent 95% confidence interval. Abbreviations: JA-Jasmonic acid, JA-Ile-Jasmonic acid isoleucine; SAG-Salicylic acid glucoside ; ABA-Abscisic acid; LOD-limit of detection.

### Change in isoprene emission, metabolic, and phytohormone levels in mechanically wounded leaves

Part of the damage done to leaves by insect chewing is mechanical damage to leaf tissue. To evaluate the impact of mechanical wounding on isoprene emission, IE leaves were wounded with forceps once photosynthesis and isoprene emission from the leaves were stable. Isoprene emission started rising within 1 min of wounding and continued to increase steadily until it became stable after 30 min (Fig. 5A).

**Fig 5.**
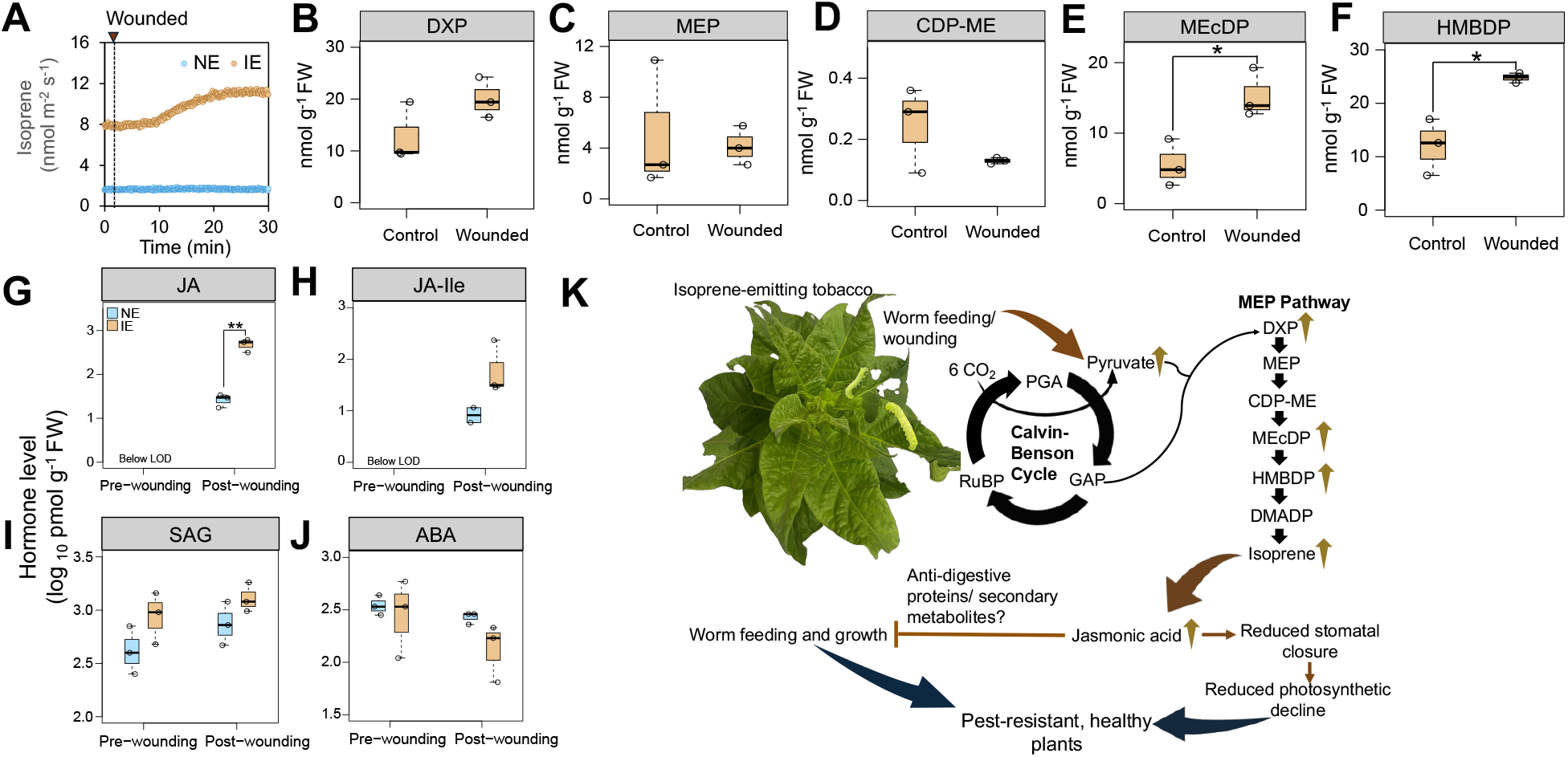
Effect of mechanical wounding on isoprene emission, MEP pathway metabolites, and hormones. **(A)** Change in isoprene emission in NE and IE leaves after wounding. **(B-F)** Change in MEP pathway metabolites in wounded IE leaves compared to unwounded (control) leaves. Increase in HMBDP and MEcDP is significantly higher after wounding (*P*<0.05; Student’s t-test). **(G-J)** Change in hormone levels in NE and IE leaves after mechanical wounding. Hormones were quantified in leaves 1 h post-wounding (*n*=3). Asterisks indicate significantly higher JA levels in IE leaves post-wounding compared to NE leaves (*P*<0.01; Student’s t-test). Whiskers of the box plots represent 95% confidence interval. **(K)** Mechanism of isoprene-mediated pest resistance. Increased JA levels in IE plants may lead to elevated levels of defense compounds making IE plants more resistant to insect herbivory. Abbreviations: DXP-1-deoxy-D-xylulose-5-phosphate; MEP-methylerythritol 4-phosphate; CDP-ME-4-(cytidine-5’-diphospho)-2-C-methyl-D-erythritol; MEcDP-2-C-methyl-D-erythritol-2,4-cyclodiphosphate; HMBDP-4-hydroxy-3-methylbut-2-enyl-diphosphate; JA-Jasmonic acid; JA-Ile-Jasmonic acid isoleucine; SAG-Salicylic acid glucoside ; ABA-Abscisic acid; PGA-3-phosphoglycerate; GAP-glyceraldehyde 3-phosphate; RuBP-Ribulose 1,5-bisphosphate.

To explore the underlying metabolic changes responsible for wound-induced increase in isoprene emission, levels of MEP pathway metabolites were quantified in both wounded and unwounded (control) IE leaves (Fig. 5B-F). Wounded leaves showed a significant increase in C-methyl-D-erythritol-2,4-cyclodiphosphate (MEcDP) and 4-hydroxy-3-methylbut-2-enyl-diphosphate (HMBDP) levels compared to the control (Fig. 5E, F). DXP (1-deoxy-D-xylulose-5-phosphate) displayed a similar trend, although the increase was not statistically significant (*P* = 0.15; Fig. 5B). No significant changes were observed in MEP and CDP-ME (4-cytidine-5’-diphospho-2-C-methyl D erythritol) levels after wounding (Fig. 5D, E). While DMADP peaks were not detectable by LC-MS/MS in this experiment, significant increases in the downstream metabolites like HMBDP and MEcDP presumably contributed to the elevated isoprene emission in response to wounding.

Given the observed trend of increased DXP in the wounded samples, we further analyzed post-wounding changes in CBC metabolite levels to assess if there was an enhanced substrate supply from the CBC to the MEP pathway. The level of pyruvate, a key substrate for the MEP pathway, was significantly elevated in wounded IE leaves compared to the control (Fig. S4). This change was not observed in the NE leaves. Other CBC metabolites that showed increases specifically in IE leaves after wounding included glucose 6-phosphate/ fructose 6-phosphate (G6P/ F6P), ribose 5-phosphate/ ribulose 5-phosphate/ xylulose 5-phosphate (R5P/ Ru5P/ Xu5P), and 6-phosphogluconate (6PG) (Fig. S4).

To assess whether the hormone response was similar between leaves damaged by insect feeding and mechanical wounding, the levels of JA, JA-Ile, SA, SAG, and ABA were quantified in leaf samples collected 1 h after wounding. The hormone profiles in the wounded samples resembled the pattern of insect-fed leaves, with JA showing a significant increase in IE leaves compared to NE leaves (Fig. 5G). JA-Ile exhibited a similar trend, although the increase in IE leaves was not statistically significant at the 0.05 threshold (Fig. 5H). SA was undetectable in all samples, while SAG and ABA levels did not differ significantly between NE and IE leaves before or after wounding (Fig. 5I, J).

## Discussion

Plant-derived VOCs can help mitigate pest damage either directly by producing toxic, anti-nutritional, and repellent compounds or indirectly by attracting natural predators (*32*). The interaction between plants and insects involves complex signaling mechanisms mediated by various signaling molecules including phytohormones. JA is a primary defense-related phytohormone that triggers plant immunity against herbivory via a multi-step signaling cascade (*25*). Insect herbivory induced a stronger increase in JA levels in IE plants than in NE plants (Fig. 4A). This indicates that isoprene activates downstream defense signaling pathways by priming the JA response during pest infestations in IE plants (Fig. 5K).

Plants release up to 2% of the assimilated carbon as isoprene, and in extreme environmental conditions like heat stress (e.g., 42°C), the carbon used for isoprene synthesis can surpass 25% (*21, 33*). This substantial allocation of resources towards isoprene biosynthesis suggests that isoprene offers important benefits. This study demonstrates that isoprene plays an important role in eliciting plant defense responses against pest infestations. Isoprene effectively deterred hornworm feeding (Fig. 2A-C). Isoprene has also been shown to deter parasitic wasps in a tritrophic response (*34*). However, in that case, isoprene interfered with parasitic wasps that would protect the plant, which could result in more damage of IE plants rather than less damage.

The deterrent effect of isoprene towards worm feeding was further supported by an *in vivo* feeding preference study which demonstrated that hornworms avoided feeding on IE leaves (Fig. 2D-E, Movie S1), consistent with the findings of Laothawornkitkul *et al*. (2008) (*31*). The present study also implies that isoprene may not serve as a strong signal for plant-to-plant communication, as there was no observed influence of IE plants on NE plants growing in the same pot in resisting whitefly infestation (Fig. 1) nor when IE and NE leaves were in direct contact during hornworm growth (Fig. S1). This could be due to diluted level of isoprene in the air being insufficient to activate defense responses in neighboring NE plants.

Physiological responses of plants to insect herbivory result in a more substantial reduction of photosynthetic capacity in undamaged leaf tissue than the direct loss of photosynthetic tissue to chewing damage (*35*). The decline in assimilation rates in the unwounded leaf area was less pronounced in the IE tobacco plants after worm feeding (Fig. 3A,D), similar to the response of aphid-resistant barley and wheat cultivars (*36, 37*). We further investigated the underlying mechanism of this differential photosynthetic response. Decrease in rubisco activation or RuBP regeneration sometimes leads to reduced photosynthesis following pest attack (*37*). However, changes in these parameters between IE and NE tobacco plants remained indistinguishable during worm feeding (Fig. 3G, Table S1). Rather, reduced stomatal closure in response to insect herbivory contributed to a smaller decline in photosynthesis in the IE plants.

A strong correlation between reduced stomatal conductance and decline in photosynthesis was observed in tobacco leaves exposed to hornworm oral secretions (*38*). The reduced stomatal closure may be associated with increased JA levels in IE plants, since application of coronatine (a JA-Ile analog) has been shown to delay stomatal closure (*39*). Additionally, the complete loss of photosynthetic responses in *aoc* (*allene oxide cyclase*) mutants to insect oral secretions highlights the role of JAs in regulating stomatal conductance-mediated carbon assimilation (*38*). Intracellular signaling molecules like H_2_O_2_ play important roles in guard cell signaling to promote stomatal closure (*40*). H_2_O_2_ levels are lower in IE plants compared to NE tobacco plants under unstressed conditions (Bianca M. Serda and Thomas D. Sharkey, unpublished data), further supporting the trait of reduced stomatal closure in IE plants.

Indeed, transcriptome profiling revealed that isoprene upregulates the expression of JA-biosynthetic genes including *lipoxygenases* and *12-oxophytodienoate reductase 3* (*OPR3*) in unstressed IE transgenic *Arabidopsis* and tobacco plants (*22*). Additionally, 80-100% of genes upregulated by isoprene treatment contain *cis*-regulatory elements like MYC recognition site, E-box, W-box, and GATA box, which interact with various transcription factors involved in JA signaling (*29*). Furthermore, worm feeding in tobacco plants results in jasmonate-induced accumulation of secondary metabolites such as nicotine, caffeoylputrescine, and 17-hydroxygeranyllinalool diterpenoid glycosides (HGL-DTGs) and increased levels of anti-digestive proteins like polyphenol oxidase, trypsin proteinase inhibitors, and threonine deaminase (*41*). These results together with the current data suggest that IE plants were more resistant to insect herbivory because of stronger JA-mediated signaling, leading to accumulation of insect-growth inhibitory metabolites.

On the other hand, exogenous JA affects isoprene emission, although the response varies between species. JA treatment was found to increase both local and systemic isoprene emission from *Populus tremuloides* due to an increase in newly fixed carbon (*42*). However, isoprene emission declined in *Ficus septica* after JA spraying, which correlated with reduced *ISPS* gene expression and protein levels (*43*). It will be interesting to investigate how JA signaling modulates isoprene signaling using JA-response mutants such as *jasmonic acid-insensitive* (*jai1-1*) that are prone to herbivore attack (*28*). JA also plays a significant role in the growth-defense trade-off in plants. Degradation of JASMONATE ZIM-DOMAIN (JAZ) proteins in the presence of JA subsequently activates MYC transcription factors which negatively regulate leaf growth (*44, 45*). Indeed, IE plants exhibit stunted growth relative to NE plants, suggesting a connection of isoprene in growth-defense tradeoff. It is likely that isoprene-induced higher JA levels in IE plants might activate JAZ-MYC regulatory network leading to the repression of growth-related pathways (*22*). However, the exact regulatory point of JA and isoprene interaction in mediating growth-defense tradeoff is still unclear, warranting further research at the genetic level.

The emission of various VOCs such as monoterpenes, sesquiterpenes, methyl salicylate, methyl jasmonate, and green leaf volatiles from plants typically increases in response to insect herbivory (*46*). However, isoprene emission from oak, a natural emitter, declined when the leaves experienced 40-50% chewing damage (*47*). Since the change in isoprene emission from IE tobacco plants upon short-term insect herbivory was unnoticeable, mechanical wounding was conducted to elicit a stronger response. Indeed, enhanced isoprene emission was detected from the undamaged part of the wounded IE leaves (Fig. 5A), suggesting its stress-responsive nature.

Unlike the transient isoprene burst detected in *Phragmites australis* after leaf cutting (*47*), IE tobacco leaves showed a sustained isoprene emission for at least 20 min post-wounding (Fig. 5A). The mechanism of this persistent isoprene emission was revealed from the profiling of MEP pathway and photosynthesis-related metabolites. The elevated levels of several MEP pathway metabolites in wounded IE leaves coincided with the enhanced isoprene emission upon wounding (Fig. 5B,E,F). Furthermore, the increase in pyruvate levels in the wounded leaves (Fig. S4) indicates an enhanced substrate supply from the CBC to the MEP pathway in response to wounding. This change, however, was not observed in NE plants, supporting the notion that the increased defense response of IE plants against herbivory resulted from a coordinated metabolic shift to produce more isoprene. Furthermore, post-wounding increases in 6PG and pentoses (Fig. S4) indicate an increased rate of the oxidative pentose phosphate pathway, which is crucial for supplying reducing power and intermediates important for mounting defense response against stresses including insect herbivory (*48*).

In summary, this study establishes the role of isoprene in protecting plants from pest infestations and reveals the connection of isoprene with JA response during insect herbivory. Given its deterring effect, the isoprene emission trait can be genetically introduced into non-emitting crop plants to combat insect herbivory. This isoprene emission from typically non-emitting plants may increase ozone and aerosol formation in the atmosphere. However, the magnitude of isoprene emission and whether it is sufficient to impact the local or global air quality remains to be determined. Nevertheless, enabling plants to naturally defend against pests could reduce reliance on chemical pesticides, fostering sustainable agricultural practices. Additionally, minimizing crop loss due to insect herbivory will ensure economic stability for the farmers. Thus, harnessing the protective roles of isoprene may offer a viable strategy for pest management in plants amidst climate instability.

## Acknowledgements

We thank Dr. Tony Schilmiller for his assistance with the LC-MS experiment. We also thank Cody Keilen (Growth Chamber Facility, Michigan State University), Mclaine Smith, and Erin Rhodea for their assistance with growth and maintenance of plants and Claudia Vickers for access to her IE and NE lines.

## Funding

National Science Foundation IOS-2022495 (TDS)

MSU Plant Resilience Institute

Michigan AgBioResearch

## Author contributions

Conceptualization: AS, MGM, TDS

Investigation: AS, MGM, YX

Maintenance of plants and assistance with phytohormone experiment: BMS

Phytohormone analysis methodology: JOK

Writing-original draft: AS

Writing-review and editing: AS, MGM, YX, BMS, TDS

## Data availability

All data will be deposited in Dryad (DOI provided once available).

