## Supplemental material for "Isoprene deters insect herbivory by priming plant hormone responses"

***To whom correspondence may be addressed**:

**ORCID ID:** 0000-0001-9363-6929 (AS), 0000-0002-4259-2361 (MGM), 0000-0002-8259-2315 (YX), 0009-0006-6065-6222 (BMS), 0000-0002-4423-3223 (TDS)

**The PDF file includes:**

Materials and Methods

Figs. S1 to S4

Table S1

**Other Supplementary Materials for this manuscript include the following:**

Movie S1

**Materials and Methods**

**Plant growth**

Isoprene-emitting transgenic tobacco (*Nicotiana tabacum, ‘*Samsun’) line and the non-emitting (NE) control were obtained from Claudia Vickers, University of Queensland (*17*). The NE and IE plants used in this study correspond to line 32, previously identified as the highest isoprene producer among the IE tobacco lines (*17*). The information about plasmid design, vector construction, transformation, and selection of the lines are described in Vickers *et al*. (2009) (*17*). The IE and NE lines were grown in separate greenhouses in the Michigan State University Plant Research Laboratory greenhouses. Each greenhouse had these conditions; 16-h photoperiod at a light intensity of 400-500 μmol m^-2^ s^-1^, day/night temperatures of 25-27°C/20-22°C, and 60%-65% humidity. The plants were grown from seeds in Suremix growing medium (Michigan Grower Products, Galesburg, Michigan, USA). Individual seeds were first sown onto separate trays. 14 d after germination, seedlings were transplanted into small 3.5 L pots to ensure seedling survival (five seedlings/pot). 14 d after transplantation when the seedlings are stable, they were transferred to large 7 L pots (one plant/pot).  The plants were watered with deionized water for 2 d and one-half-strength Hoagland nutrient solution for 5 d (*49*). Plants were used for experiments when they were 6 to 8-w-old, before they developed flowers and seeds.

**Insect growth and feeding assays**

Tobacco hornworm eggs (Carolina Biological Supply, Burlington, North Carolina, USA) were incubated for 2 d in a glass box with detached tobacco leaves at room temperature. After the eggs hatched, the 1^st^ instar larvae were transferred to 6-w-old plants with a paint brush. The plants were then kept in a growth chamber (16 h photoperiod, mean light intensity 400-500 μmol m^-2^ s^-1^, day/night temperatures of 25-27°C/20-22°C). Larval weights were measured using an analytical balance and consumed leaf areas were analyzed using Fiji (*50*).

**Phytohormone extraction and measurement by Liquid Chromatography-Mass Spectrometry (LC-MS/MS)**

For the short-term feeding assay, 3^rd^ instar larvae were placed on NE and IE leaves of 6-w-old plants. After 1 h of feeding, leaf discs were collected using a freeze clamp. For the long-term feeding assay, 1^st^ instar larvae were placed on NE and IE leaves of 6-w-old plants and allowed to feed for 10 d. For mechanical wounding, a part of the leaf outside the gas exchange chamber was wounded with forceps and leaf discs within the gas exchange chamber were collected after 1 h. Frozen leaf discs were ground into a fine powder using a tissue homogenizer (Mixer Mill MM301, Retsch, Newton, PA, USA). Then 800 μL of ice-cold extraction buffer (80:20 v/v methanol: water, 0.1% formic acid, 0.1 g/L butylated hydroxytoluene) was added to the ground plant material. 100 nM of labeled internal standards including SA-^13^C_6_ (Santa Cruz Biotechnology, sc-220088), ABA-d_6_ (Toronto Research Chemicals, A110002), and JA-d_5_ (Cayman Chemical, 29076) were added to the extraction buffer. The tubes were vortexed and placed on a rocking platform at 4°C for 24 h. On the next day, the samples were vortexed to mix and then centrifuged at 12,000 g for 10 min at 4°C. Then 400 μL supernatant was transferred to centrifugal filter units (PALL Nanosep with 0.2µm polytetrafluoroethylene membrane) and centrifuged at 5,000 g for 1 min at room temperature. The flow-through was collected and aliquoted into 2 mL glass vials with inserts for LC-MS/MS analysis. Samples were analyzed by HPLC after extraction. The hormones were separated using the Acquity UPLC BEH C18 column (2.1 x 50 mm, 1.7 micron) fitted on a Xevo TQ-XS mass spectrometer. The chromatographic separation utilized a multi-step gradient with mobile phase A (water + 0.1% formic acid) and mobile phase B (acetonitrile): 0-0.5 min, 95% A; 0.5-10 min, 95-70% A; 10-11 min, 70-5% A; 11-13 min, 5% A; 13-13.01 min, 5-95% A, 13.01-15 min, 95% A, at a flow rate of 0.5 mL min^−1^. Column temperature was maintained at 40°C. The source temperature was maintained at 150°C, and the desolvation temperature was set to 400°C.

**Gas exchange measurements**

Measurements of photosynthesis (*A*), intercellular CO₂ concentration (*C_i_*), and stomatal conductance (*g_sw_*) in NE and IE leaves were taken using a LI-COR 6800 portable photosynthesis system (LI-COR Biosciences, Lincoln, NE, USA). A fully expanded mature leaf was clamped to a 6 cm^2^ chamber of LI-COR 6800 and maintained under CO₂ concentration of 420 µmol mol⁻¹, light intensity of 1,000 μmol m⁻² s⁻¹, a temperature of 28°C, and a water vapor pressure difference (VPD) of 1.3 kPa. Photosynthesis was allowed to stabilize which took between 30 min and 1 h. Once photosynthesis was stable, three to five 3^rd^ instar hornworm larvae were placed on a part of the leaf outside the chamber and photosynthetic measurements were recorded every 5 seconds for 45 min during worm feeding.

***A/Ci* curve measurements**

Representative *A/Ci* curves were generated using the dynamic assimilation technique in both NE and IE leaves during worm feeding. These curves were obtained using a LI-COR 6800 portable photosynthesis system (LI-COR Biosciences, Lincoln, NE, USA) under a light intensity of 1,000 μmol photons m⁻² s⁻¹, a leaf temperature of 28°C, and a VPD of 1.3 kPa. The sequence of reference CO₂ concentrations used was 50, 100, 200, 300, 350, 400, 450, 500, 550, 600, 700, 800, 1,000, 1,200, and 1,500 µmol mol⁻¹, as outlined by Sharkey (2019) (*51*). *A/Ci* from four replicates were fitted using the routine described by Sharkey (2015)(*52*). Photosynthesis and *A/Ci* curve parameters including the rubisco carboxylation capacity (*V_cmax_*), the electron transport capacity (*J*), the triose phosphate utilization (*TPU*) capacity, respiration in the light (*R_L_*), and mesophyll conductance to CO_2_ transfer (*g_m_*), were estimated using the method described by Sharkey (2015) (*52*). Additionally, the proportion of carbon exported from photorespiration as glycine (*α_g_*) or serine (*α_s_*) was estimated using equations described by Busch *et al*. (2018) (*53*).

**Isoprene measurement in wounded leaves**

Isoprene emission measurements were conducted using a Fast Isoprene Sensor (FIS; Hills Scientific, Boulder, Colorado). A fully expanded mature leaf of IE plant was clamped to a 6 cm^2^ chamber of LI-COR 6800. The leaf was allowed to equilibrate under the following conditions: light intensity of 1000 µmol m^-2^ s^-1^ (50% blue light and 50% red light), temperature of 30°C, CO_2_ of 420 µmol mol^-1^ and water vapor content of 22-26 mmol mol^-1^ depending on laboratory room temperature. Then a part of the leaf outside the chamber was wounded with forceps. Exhaust air from the LI-COR 6800 was fed into the FIS for isoprene measurements. The flow rate in the LI-6800 was set at 500 μmol s^-1^ and the FIS flow rate was set such that it drew sample air from the LI-6800 at 420 μmol s^-1^. A 3.225 ppm isoprene standard was used for the FIS calibration. Isoprene emission measurements were logged every 5 s.

**MEP pathway** **metabolites extraction and measurement by LC-MS/MS**

Leaf discs were collected using a freeze clamp 20 min after wounding with forceps. MEP pathway metabolites were extracted using the protocol described by Sahu *et al*. (2023)(*54*). Frozen leaf discs were ground into a fine powder using a tissue homogenizer (Mixer Mill MM301, Retsch, Newton, PA, USA) and extracted for 15 min using 300 μL of ice-cold extraction buffer containing 3:1:1 acetonitrile: isopropanol: 20 mM ammonium bicarbonate, adjusted to pH 10 with ammonium hydroxide. The samples were then centrifuged at 14,000 g for 10 min. The supernatant was transferred to 2 mL glass vials with glass inserts and analyzed by LC-MS/MS immediately following the extraction. Standards of the MEP pathway metabolites including DXP, MEP, CDP-ME, MEcDP, and HMBDP (Echelon Biosciences, Logan, UT, USA) were separated using InfinityLab Poroshell 120 HILIC-Z, P column (2.1 x 100 mm, 2.7 micron with column ID) fitted on a Xevo TQ-XS mass spectrometer. Column temperature was maintained at 25°C. The mobile phase consisted of 20 mM ammonium bicarbonate adjusted to pH 10.0 with ammonium hydroxide and acetonitrile. Negative mode electrospray ionization was used with the following settings: capillary 1.00 kV, source temperature of 150°C, and desolvation temperature of 400°C.

**CBC** **metabolites extraction and measurement by Ion-Pair Chromatography – Tandem Mass Spectrometry (IPC-MS/MS)**

For the analysis of phosphorylated intermediates in the Calvin-Benson Cycle (CBC), metabolites were extracted from rapidly frozen tissues using the protocol previously described by Xu *et al*. (2021) (*55*). Mass spectrometry analyses were performed following the methods outlined in earlier studies (*56, 57*). Ion-pair chromatography – tandem mass spectrometry (IPC-MS/MS) was performed using an Acquity UPLC pump system (Waters, Milford, MA, USA) coupled with a Waters Xevo TQ-S UPLC/MS/MS (Waters, Milford, MA, USA). Metabolites were separated on a 2.1×50 mm Acquity UPLC BEH C18 Column (Waters, Milford, MA, USA) at 40°C. The chromatographic separation utilized a multi-step gradient with mobile phase A (10 mM tributylamine in 5%(v/v) methanol) and mobile phase B (methanol): 0-1 min, 95-85% A; 1-6 min, 65-40% A; 6-7 min, 40-0% A; 7-8 min, 0% A; 8-9 min, 100% A, at a flow rate of 0.3 mL min^−1^. The source temperature was maintained at 120°C, and the desolvation temperature was set to 350°C. Nitrogen served as the sheath and auxiliary gas, with collision gas (argon) set to 1.1 mTorr. Gas flow for desolvation and cone was adjusted to 800 and 50 L/h, respectively. The scan time was 0.1 ms.

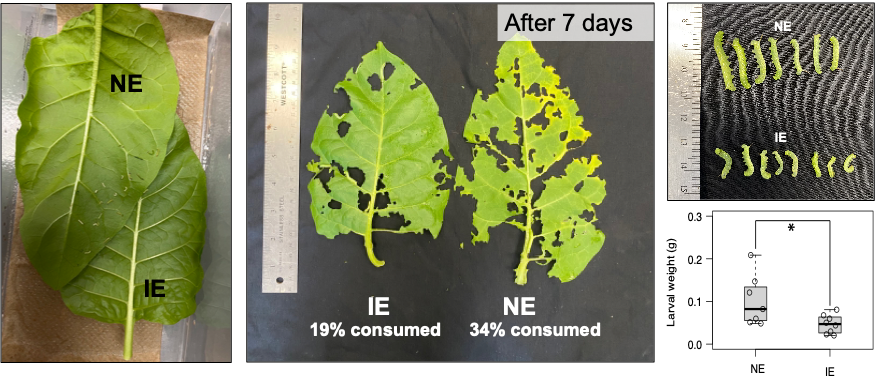

**C**

**B**

**A**

**Fig. S1**. **Hornworm feeding preference study after hatching.** **(A)** Worms were reared in a box containing a pair of IE and NE leaves for 7 d. **(B)** Comparison of leaf consumption 7 d post-feeding when given the choice between NE and IE leaves. **(C)** Comparison of hornworm larval weight recovered from IE and NE leaves after 7 d (*n*=7-8). Asterisk indicates significantly lower larval weight in IE leaves compared to NE leaves 10 d post-feeding (*P*<0.05; Student’s t-test).

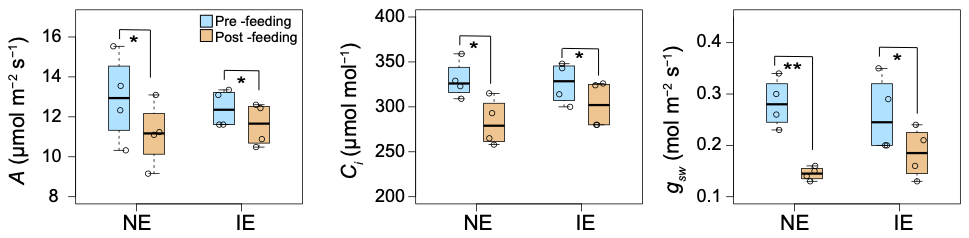

**C**

**B**

**A**

**Fig. S2**. **Comparison of (A) photosynthesis (*A*), (B) intercellular CO_2_ concentration (*C_i_*), and (C) stomatal conductance (*g_sw_*) in NE and IE leaves pre- and post-feeding.** Asterisks indicate significant decrease after 45 min worm feeding (*- *P*<0.01; **- *P*<0.01; Student’s t-test). Whiskers of the box plots represent 95% confidence interval.

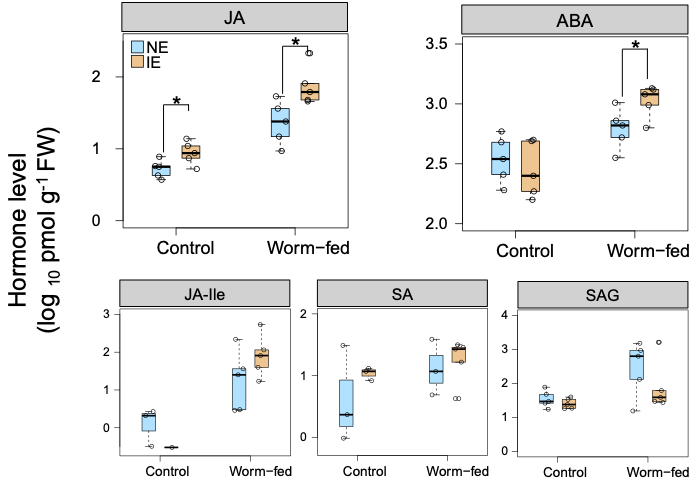

**Fig. S3**. **Endogenous hormone levels in NE and IE leaves after long-term worm feeding.** Hormones were quantified in leaves 10 d post-feeding (*n*=3-5). Asterisk indicates significantly higher JA and ABA levels in IE leaves post-feeding compared with NE leaves (*P*<0.05; Student’s t-test). Change in JA-Ile, SA, and SAG levels between NE and IE lines was not statistically significant. Control plants were never exposed to worms. Whiskers of the box plots represent 95% confidence interval. Abbreviation: SA- Salicylic acid.

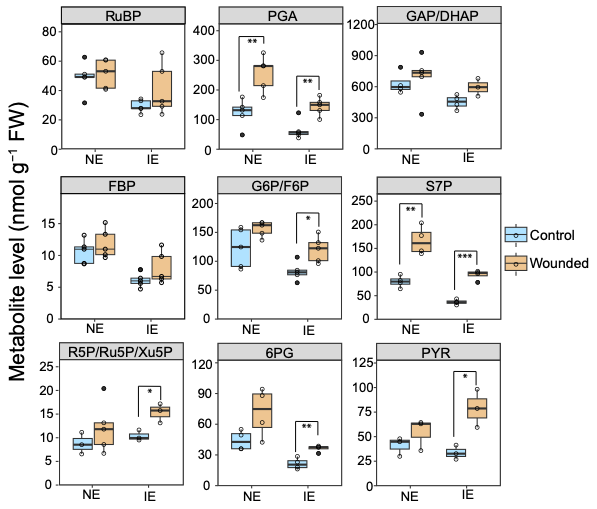

**Fig. S4**. **Change in CBC metabolites after wounding in NE and IE leaves.** Asterisks indicate significant increase in wounded leaves compared to unwounded (control) leaves (*- *P*<0.01; **- *P*<0.01; ***- *P*<0.001; Student’s t-test). Whiskers of the box plots represent 95% confidence interval. Abbreviations: FBP- fructose 6(bis)-phosphate; RuBP- Ribulose 1,5-bisphosphate; PGA- 3-phosphoglycerate; G6P- glucose 6-phosphate; F6P- fructose 6-phosphate; R5P- ribose 5-phosphate; Ru5P, ribulose 5-phosphate; Xu5P- xylulose 5-phosphate; 6PG- 6 phosphogluconate; S7P- sedoheptulose 7(bis)-phosphate; PYR- pyruvate; GAP- glyceraldehyde 3-phosphate; DHAP- dihydroxyacetone phosphate.

|  | **Units** | **NE** | **IE** |
| --- | --- | --- | --- |
| *V_cmax_* | μmol m^-2^ s^-1^ | 73.5 ± 4.9 | 77.7 ± 9.9 |
| *J* | μmol m^-2^ s^-1^ | 116.1 ± 10.9 | 114.0 ± 4.6 |
| *TPU* | μmol m^-2^ s^-1^ | 7.8 ± 0.9 | 7.4 ± 0.4 |
| *R_d_** | μmol m^-2^ s^-1^ | 2.0 ± 0.0 | 2.0 ± 0.0 |
| *g_m_* | μmol m^-2^ s^-1^Pa^-1^ | 1.5 ± 0.3 | 0.9 ± 0.2 |
| *α_G_* | none | 0.0 ± 0.0 | 0.0 ± 0.0 |
| *α_S_* | none | 0.3 ± 0.1 | 0.4 ± 0.0 |

**Table S1: Photosynthesis and *A/Ci* curve parameters in NE and IE leaves recorded during worm feeding.** No significant difference was observed between the NE and IE lines. *V_cmax_*– Rubisco capacity; *J*- electron transport; *TPU*- triosephosphate use; *R_d_*- respiration; *g_m_*- mesophyll conductance; *α_G_*- proportion of carbon exported from photorespiration as glycine; *α_S_*- proportion of carbon exported from photorespiration as serine.

**Movie S1: Hornworm feeding preference study.** Feeding behavior of 3^rd^ instar larvae was monitored when given the choice between NE and IE leaves. Although the worms crawled towards the IE leaf, they turned away and preferred to feed on the NE leaf.
